# Large-scale sequence comparisons with *sourmash*

**DOI:** 10.1101/687285

**Authors:** N. Tessa Pierce, Luiz Irber, Taylor Reiter, Phillip Brooks, C. Titus Brown

## Abstract

The sourmash software package uses MinHash-based sketching to create “signatures”, compressed representations of DNA, RNA, and protein sequences, that can be stored, searched, explored, and taxonomically annotated. sourmash signatures can be used to estimate sequence similarity between very large data sets quickly and in low memory, and can be used to search large databases of genomes for matches to query genomes and metagenomes. sourmash is implemented in C++, Rust, and Python, and is freely available under the BSD license at http://github.com/dib-lab/sourmash.

## Introduction

Bioinformatic analyses rely on sequence comparison for many applications, including variant analysis, taxonomic classification and functional annotation. As the Sequence Read Archive now contains over 20 Petabases of data (1), there is great need for methods to quickly and efficiently conduct similarity searches on a massive scale. MinHash techniques (2) utilize random sampling of k-mer content to generate small subsets known as “sketches” such that Jaccard similarity (the intersection over the union) of two sequence data sets remains approximately equal to their true Jaccard similarity (2, 3). The many-fold size reductions gained via MinHash open the door to extremely large scale searches.

While the initial k-mer MinHash implementation focused on enabling Jaccard similarity comparisons (3), it has since been modified and extended to enable k-mer abundance comparisons (4), decrease runtime and memory requirements (5), and work on streaming input data (6). Furthermore, as Jaccard similarity is impacted by the relative size of the sets being compared, containment searches (2, 7, 8) have been developed to enable detection of a small set within a larger set, such as a genome within a metagenome.

Here we present version 2.0 of sourmash (9), a Python library for building and utilizing MinHash sketches of DNA, RNA, and protein data. sourmash incorporates and extends standard MinHash techniques for sequencing data, with a particular focus towards enabling efficient containment queries using large databases. This is accomplished with two modifications: (1) building sketches via a modulo approach (2), and (2) implementing a modified Sequence Bloom Tree (10) to enable both similarity and containment searches. In most cases, these features enable sourmash database comparisons in sub-linear time.

Standard genomic MinHash techniques, first implemented in (3), retain the minimum *n* k-mer hashes as a representative subset. sourmash extends these methods by incorporating a user-defined “scaled” factor to build sourmash sketches via a modulo approach, rather than the standard bottom-hash approach (2). Sketches built with this approach retain the same fraction, rather than number, of k-mer hashes, compressing both large and small datasets at the same rate. This enables comparisons between datasets of disparate sizes but can sacrifice some of the memory and storage benefits of standard MinHash techniques, as the signature size scales with the number of unique k-mers rather than remaining fixed (8). In sourmash, use of the “scaled” factor enables user modification of the trade-off between search precision and sketch size, with the caveat that searches and comparisons can only be conducted using signatures generated with identical “scaled” values (downsampled at the same rate).

To enable large-scale database searches using these signatures, sourmash implements a modified Sequence Bloom Tree, the SBTMH, that allows both similarity (sourmash search) and containment (sourmash gather) searches for taxonomic exploration and classification. Notably, Jaccard similarity searches using this modified SBT require storage of the cardinality of the smallest MinHash below each node in order to properly bound similarity. sourmash also implements a second database format, LCA, for in-memory search when sufficient RAM is available or database size is tractable.

In addition to these modifications, sourmash implements k-mer abundance tracking (4) within signatures to allow abundance comparisons across datasets and facilitate metagenome, metatranscriptome, and transcriptome analyses, and is compatible with streaming approaches. The sourmash library is implemented in C++, Rust (11), and Python, and can be accessed both via command line and Python API. The code is available under the BSD license at http://github.com/dib-lab/sourmash.

## Implementation

sourmash provides a user-friendly, extensible platform for MinHash signature generation and manipulation for DNA, RNA, and protein data. Sourmash is designed to facilitate containment queries for taxonomic exploration and identification while maintaining all functionality available via standard genomic MinHash techniques.

### sourmash Signatures

sourmash modifies standard genomic MinHash techniques in two ways. First, sourmash scales the number of retained hashes to better represent and compare datasets of varying size and complexity. Second, sourmash optionally tracks the abundance of each retained hash, to better represent data of metagenomic and transcriptomic origin and allow abundance comparisons.

#### Scaling

sourmash implements a method inspired by modulo sketches (2) to dynamically scale hash subset retention size (*n*). When using scaled signatures, users provide a scaling factor (*s*) that divides the hash space into *s* equal bands, retaining hashes within the minimum band as the sketch. These scaled signatures can be converted to standard bottom-hash signatures, if the subset retention size *n* is equal to or smaller than the number of hashes in the scaled signature. sourmash provides a signature utility, downsample, to convert sketches when possible. Finally, to maintain compatibility with sketches generated by other programs such as Mash (3), sourmash generates standard bottom-hash MinHash sketches if users specify the hash subset size *n* rather than scaling factor.

#### Streaming Compatibility

Scaled signature generation is streaming compatible and provides some advantages over streaming calculation using standard MinHash. As data streams in, standard MinHash replaces hashes based on the minimum hash value to maintain a fixed number of hashes in the signature. In contrast, no hash is ever removed from a scaled signature as more data is received. As a result, for searches of a database using streamed-in data, all prior matches remain valid (although their significance may change as more data is received). This allows us to place algorithmic guarantees on containment searches using streaming data.

#### Abundance tracking

sourmash extends MinHash functionality by implementing abundance tracking of k-mers. k-mer counts are incremented after hashing as each k-mer is added to the hash table. sourmash tracks abundance for k-mers in the minimum band and stores this information in the signature. These values accompany the hashes in downstream comparison processes, making signatures better representations of repetitive sequences of metagenomic and transcriptomic origin.

#### Signatures

MinHash sketches associated with a single sequence file are stored together in a “signature” file, which forms the basis of all sourmash comparisons. Signatures may include sketches generated with different *k* sizes or molecule type (nucleotide or protein) and are stored in JSON format to maintain human readability and ensure proper interoperability.

Signatures can only be compared against signatures and databases made from the same parameters (*k* size(s), scaled value, nucleotide or protein level). If signatures differ in their scaled value or size(*n*), the larger signatures can be downsampled to become comparable with smaller signatures using the *signature utilities*, below. sourmash also provides 6-frame nucleotide translation to generate protein signatures from nucleotide input if desired.

#### Signature Utilities

sourmash provides a number of utilities to facilitate set operations between signatures (merge, intersect, extract, downsample, flatten subtract, overlap), and handling (describe, rename, import, export) of sourmash signatures. These can be accessed via the sourmash signature subcommand.

### SBT-MinHash

sourmash implements a modified Sequence Bloom Tree (SBT (10)), the SBT-MinHash (SBTMH), which can efficiently capture large volumes of MinHashes (e.g., all microbes in GenBank) and support multiple search regimes that improve on run time of linear searches.

#### Implementation

The SBTMH is a n-ary tree (binary by default), where leaf nodes are MinHash signatures and internal nodes are Bloom Filters. Each Bloom Filter contains all the values from its children, so the root node contains all the values from all signatures. SBTMH is designed to be extensible such that signatures can be subsequently added without the need for full regeneration. Adding a new signature to SBTMH causes parent nodes up to the root to be updated, but other nodes are not affected.

SBTMH trees can be combined if desired: In the simplest case, adding a new root and updating it with the content of the previous roots is sufficient, and this preserves all node information without changes. As an example, separate indices can be created for each RefSeq subdivision (bacteria, archaea, fungi, etc) and be combined depending on the application (such as an analysis for bacteria + archaea, but not fungi). In practice, this is most useful for updating the SBTMH, as both search and gather support search over multiple databases without the need for rebuilding a single large database.

#### Searching SBTMH

Similarity searches start at the root of the SBTMH, and check for query elements present in each internal node. If the similarity does not reach the threshold, the subtree under that node does not need to be searched. If a leaf is reached, it is returned as a match to the query signature. In order to enable similarity (in addition to containment) searches using this modified SBTMH, nodes store the cardinality of the smallest signature below each node in order to properly bound similarity. The full SBTMH does not need to be imported to RAM during searches, making this method best for rapid searching with minimal memory requirements. However, if sufficient RAM is available, searches of databases (or many signatures) may be completed in memory via an alternate database format (discussed below).

#### SBTMH Utilities

sourmash provides several utilities for construction, use, and handling of SBTMH databases. These include sbt index to index a collection of signatures as an SBTMH for fast searching, sbt append to add signatures, and sbt combine to join two SBTMH databases.

### LCA Database

sourmash implements an alternate database format, LCA, to support in-memory queries. This implementation utilizes two named lists to store MinHashed databases: the first containing MinHashes, and the second containing taxonomic information, with both lists named by sample name. This structure facilitates direct look-up of MinHashes, and thus can be leveraged to return additional information, such as taxonomic lineage. The LCA database structure can be prepared using the sourmash lca index command.

## Assessing Sequence Similarity

### Pairwise Comparisons

For sequence comparison, sourmash compare re-implements Jaccard sequence similarity comparison to enable comparison between scaled MinHashes. When abundance tracking of k-mers is enabled, compare instead calculates the cosine similarity, although we recommend using more accurate approaches for detailed comparisons (6).

### Database Searches

In addition to conducting pairwise comparisons, two types of database searches are implemented: breadth-first similarity searches (sourmash search) and best-first containment searches (sourmash gather), which support different biological queries. These searches con be conducted using either database format.

#### Similarity Queries

Breadth-first sourmash search can be used to obtain all MinHashes in the SBTMH that are present in the query signature (above a specified threshold). This style of search is streaming-compatible, as the query MinHash can be augmented as the search is occurring.

#### Containment Queries

Best-first sourmash gather implements a greedy algorithm where the SBTMH is descended on a linear path through a set of internal nodes until the highest containment leaf is reached. The hashes in this leaf are then subtracted from the query MinHash and the process is repeated until the threshold minimum is reached. sourmash post-processes similarity statistics after the search such that it reports percent identity and unique identity for each match.

#### Taxonomy-resolved searches

sourmash can conduct taxonomy-resolved searches uses the “least common ancestor” approach (as in Kraken (12)), to identify k-mers in a query. From this it can either find a consensus taxonomy between all the k-mers (sourmash classify) or it can summarize the mixture of k-mers present in one or more signatures (sourmash summarize).

## Use Cases

Below we provide several use cases to demonstrate the utility of sourmash for sequence comparisons. We primarily demonstrate nucleotide-level applications in this paper; protein-level sourmash will be explored further in future work. Additional information and tutorials are available at https://sourmash.readthedocs.io.

### Installation

To install sourmash, we recommend using conda:

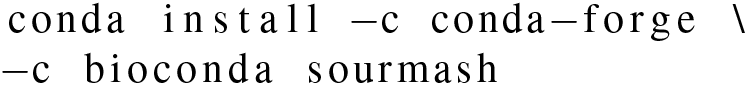

Alternate installation instructions are available at sourmash.readthedocs.io.

### Creating a signature

All sourmash comparisons work on signatures, compressed representations of biological sequencing data. To create a signature from sequences with abundance tracking:

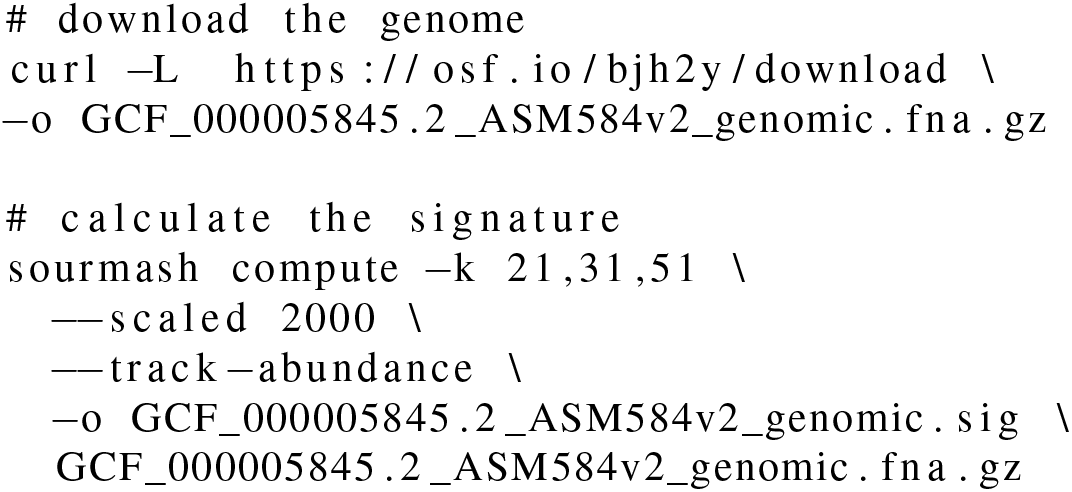

Because a signature can contain multiple MinHashes, multiple k-sizes can be specified per a sequence. Only one scaled size can be used.

By default, the name of the file becomes the name of the signature. To name the signature from the first line of the sequencing file, use ––name–from–first. Although the ––track–abundance flag is optional, since downstream comparison methods contain the flag ––ignore–abundance to ignore them, we recommend calculating all signatures with abundance tracking.

To create a signature from protein sequences:

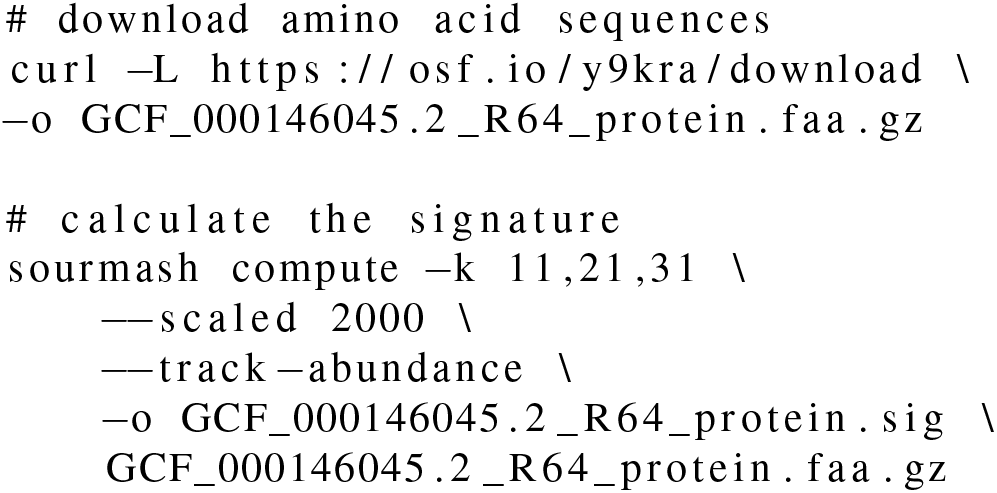

Signatures can also be made directly from reads. Depending on the downstream use cases, we recommend different preparation methods. When the user aims to compare the signature to other signatures, we recommend k-mer trimming the reads before computing the signature. Because compare does an all-by-all comparison of signatures, errors in the reads will falsely deflate the similarity metric. We recommend trimming RNA-seq or metagenome reads with trim low abund.py in the khmer package (13), a dependency of sourmash.

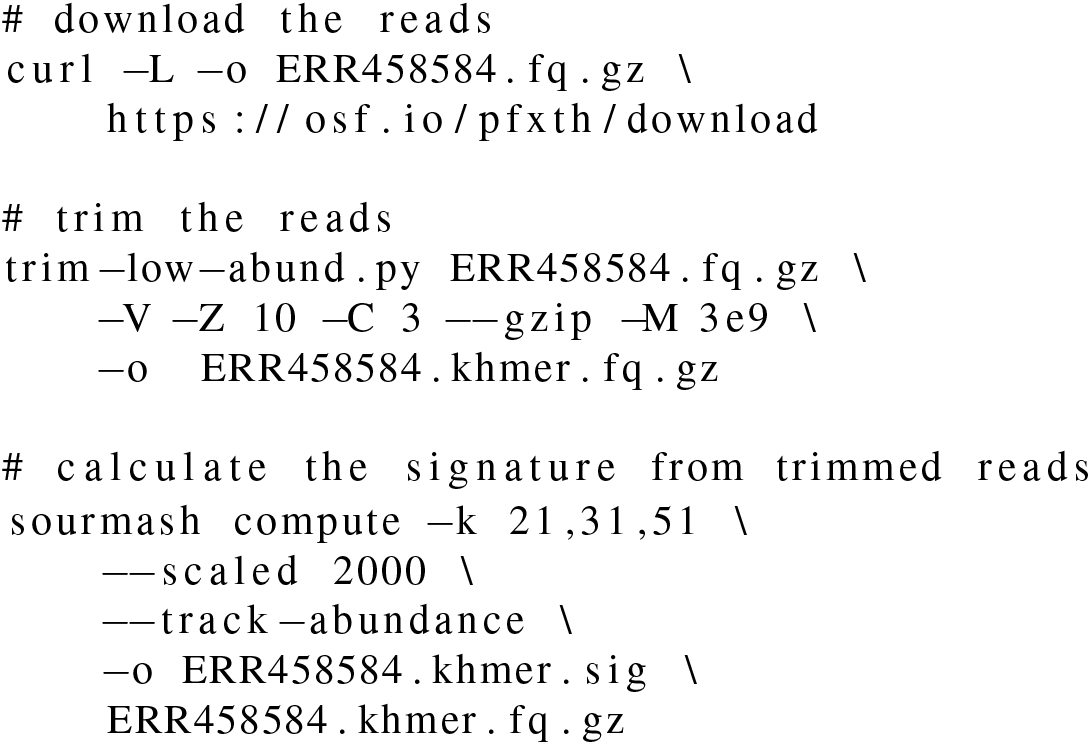

When using methods that compare a signature against a database such as gather or search, k-mer trimming need not be used. These methods use exact matching of hashes in the signature to those in the databases. k-mer trimming could increase false negatives, but results on k-mer trimmed data will more accurately represent the proportions of content in the data.

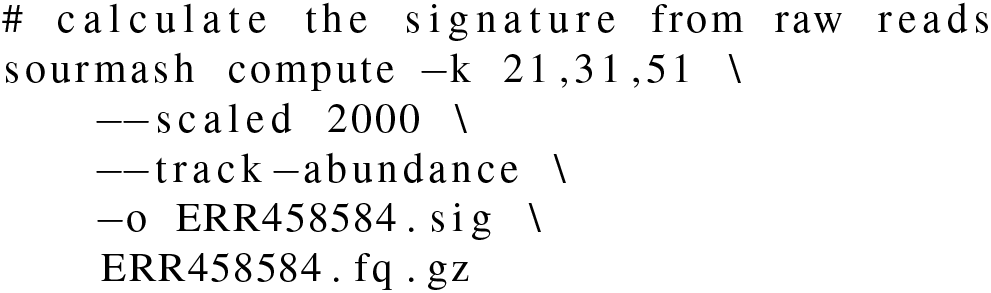

### Comparing many signatures

Signatures calculated with abundance tracking enable rapid comparison of sequences where k-mer frequency is variable, and can be leveraged for quality control and summarization methods. For example, principle component analysis (PCA) and multidimensional scaling (MDS) are standard quality control and summarization methods for count data generated during RNA-seq analysis (14). sourmash can be used to build this MDS plot in a reference-free (or assembly-free) manner, using k-mer abundances of the input reads.

### MDS

Here, we use a set of four *Saccharomyces cerevisiae* RNA-seq samples: replicate wild-type samples and replicate mutant (*SNF2*) samples (15). To use sourmash to build an MDS plot, we first trim the data to remove low abundance k-mers via khmer (13). We demonstrate the streaming capability of sourmash by downloading, k-mer trimming, and calculating a signature with one command. This allows the user to generate signatures without needing to store large files locally.

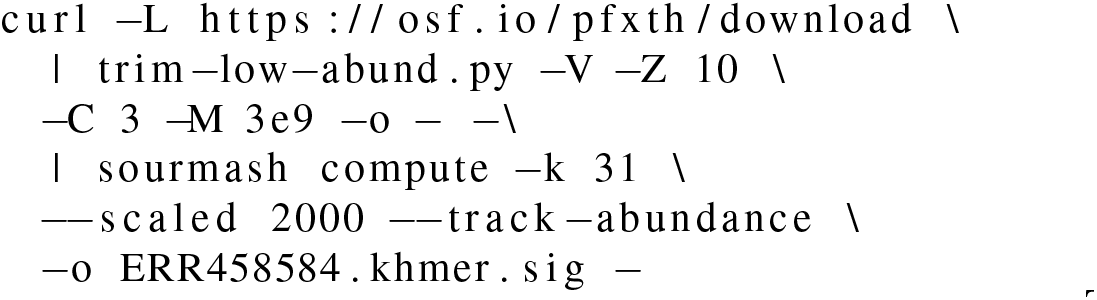

The signature will be named from the input filename, in this case . We can change the name to reflect its contents using the signature rename function.

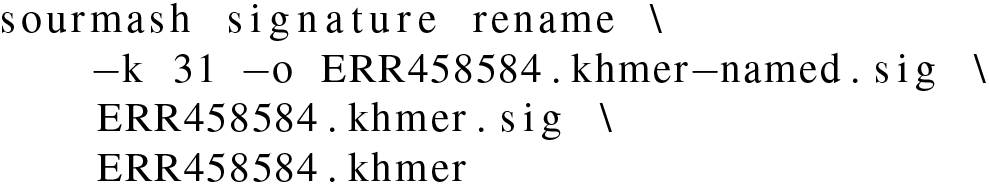

We can also check that the name has been changed.

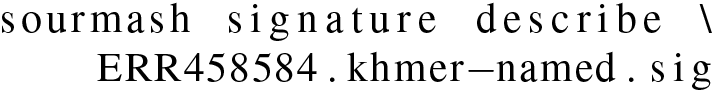

Using signatures from four samples, we can compare the files with the compare function. Here we download signatures calculated and renamed using the above commands. We output the comparison matrix as a csv for downstream use in other platforms.

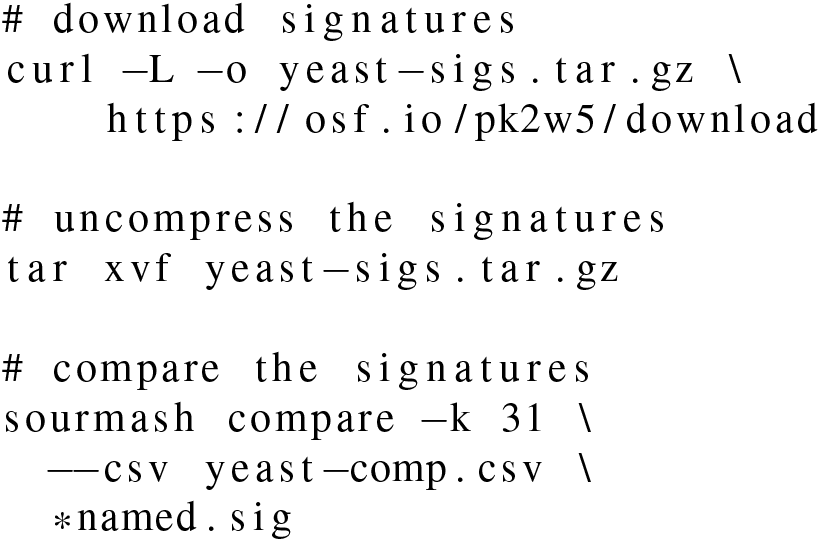

We then import the compare similarity matrix into R to produce an MDS plot with wild-type samples (ERR459011, ERR459102) in yellow and mutant samples (ERR458584, ERR458829) in blue.

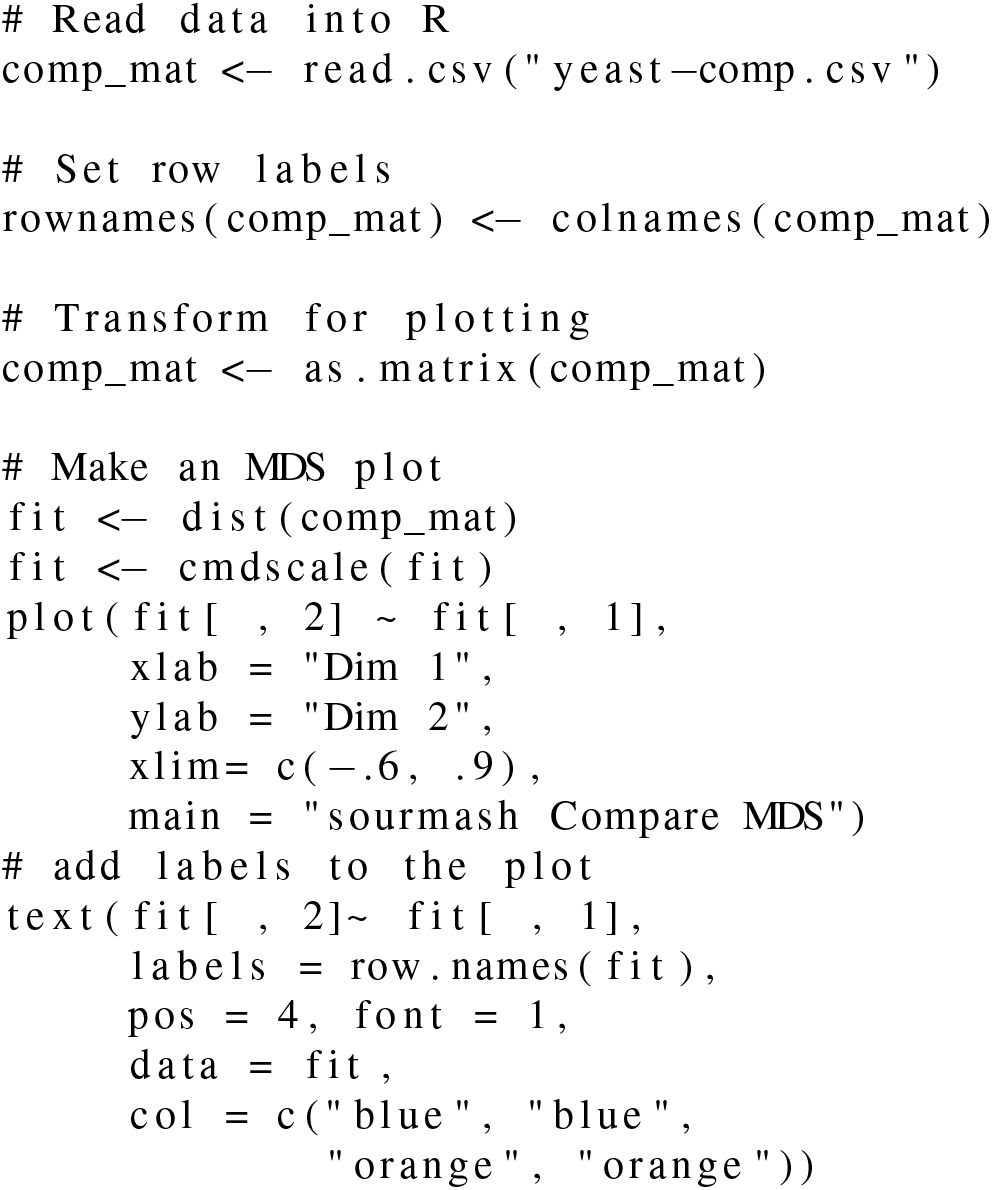

To see how the sourmash MDS plot compares with traditional methods using transcript count data, we used Salmon (16) to quantify abundance relative to an *S. cerevisiae* reference, and edgeR (17) to build an MDS plot. Comparison code and figure are available in the Appendix.

We also find this useful for comparing other types of RNA sequencing samples (mRNA, ribo-depleted, 3’ tag-seq, meta-transcriptomes, and transcriptomes).

### Tetranucleotide Frequency Clustering

We can also use sourmash with abundance tracking for tetranucleotide frequency clustering. Tetranucleotide usage is species-specific, with strongest conservation in DNA coding regions (18). This is often used in metagenomics as one method to “bin” assembled contiguous sequences together that are from the same species (19). Recently, tetranucleotide frequency clustering using sourmash was used to detect microbial contamination in the domesticated olive genome (20). Here we reimplement this approach using 100 of the 11,038 scaffolds in the draft genome. We calculate the signature using a k-mer size of 4, use all 4-mers, and track abundance. Then we use sourmash compare to calculate the similarity between each scaffold. (The ––singleton flag, calculates a signature for each sequence in the fasta file.)

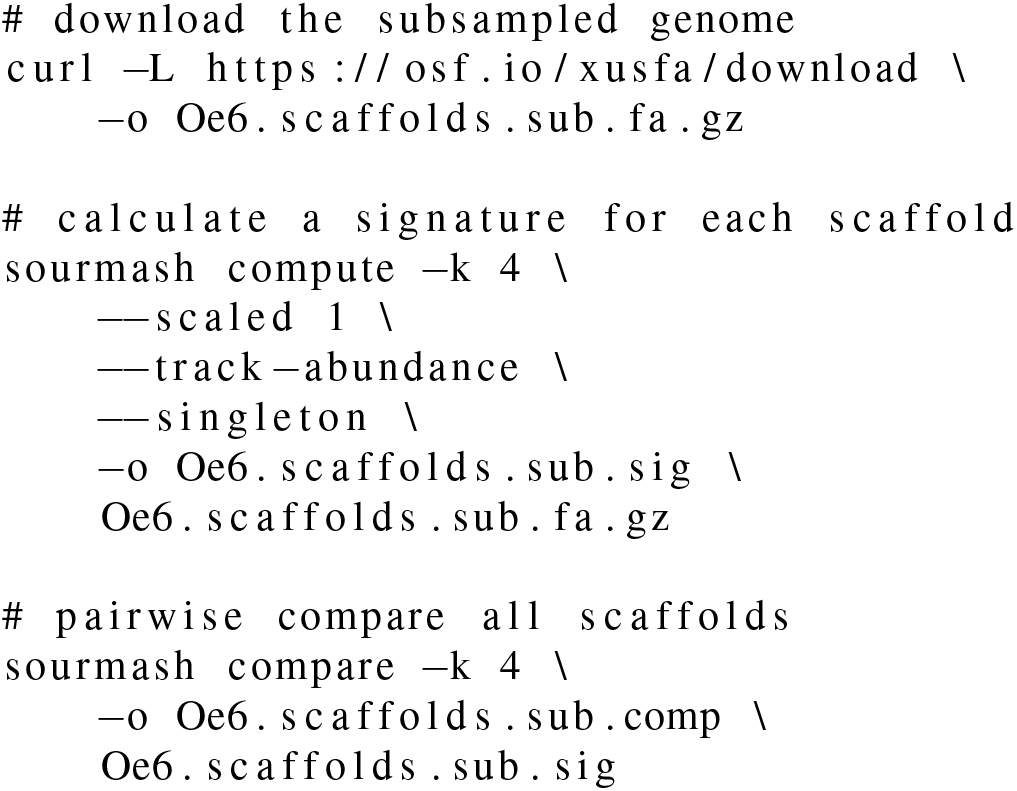

Although sourmash compare supports export to a csv file, sourmash also has a built in visualization function, plot . We will use this to visualize scaffold similarity.

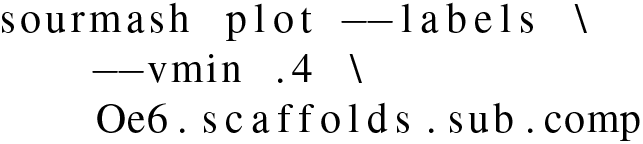

In Figure 2, we see that there is high similarity between 98 of the scaffolds, but that Oe6_s01156 and Oe6_s01003 are outliers with tetranucleotide frequency similarity around 40% to olive scaffolds. These two scaffolds are contaminants (20).

**Fig. 1.**
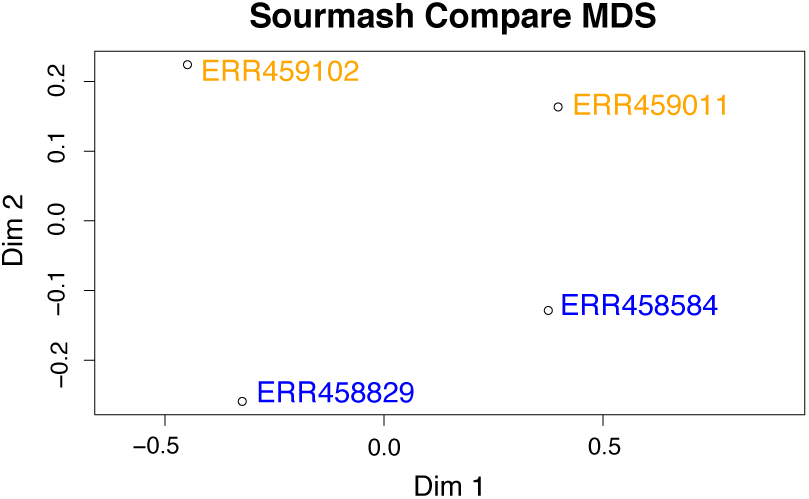
MDS plot generated using sourmash signatures built from k-mer trimmed reads. Wild-type samples (ERR459011, ERR459102) are in yellow and mutant samples (ERR458584, ERR458829) in blue.

**Fig. 2.**
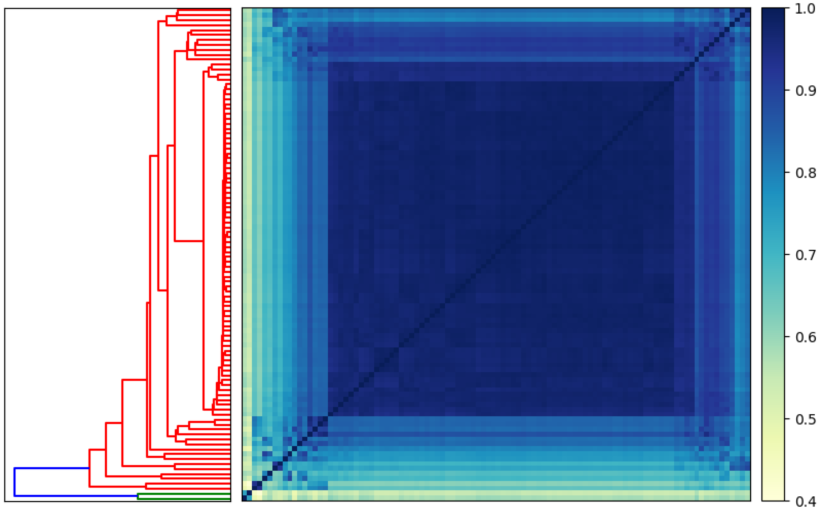
Heatmap and dendrogram generated using sourmash signatures built from scaffolds in the domesticated olive genome. Two scaffolds are outliers when using tetranucleotide frequency to calculate similarity (highlighted in green on the dendrogram).

### Comparisons to detect outliers

MinHash comparisons are useful for outlier detection. Below we compare 50 genomes that contain the word “*Escherichia coli*.” We have pre-calculated the signatures for each of these genomes. We then use the plot function to visualize our comparison.

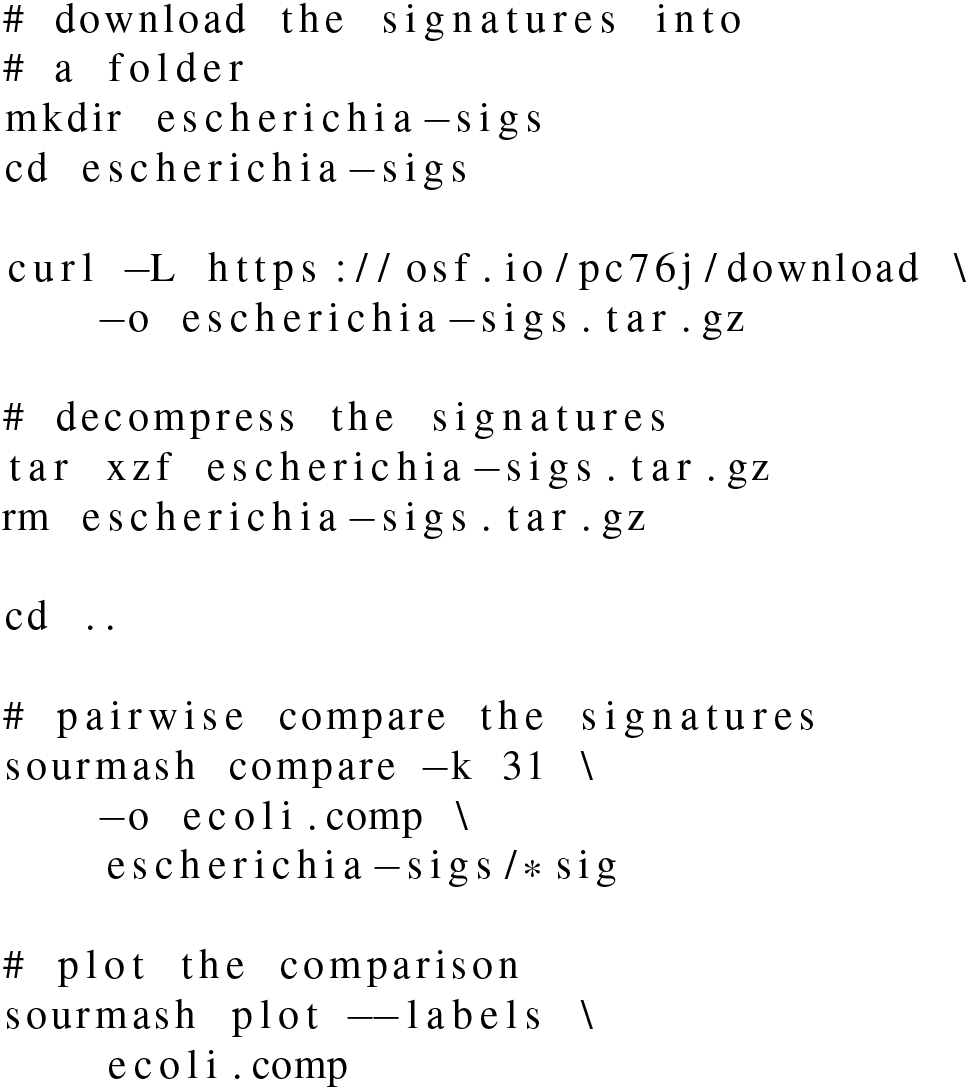

We see that the minimum similarity in the matrix is 0%. If all 50 signatures were from the same species, we would expect to observe higher minimum similarity at a k-mer size of 31. When we look closely, we see one signature has 0% similarity with all other signatures because it is a phage (Figure 3).

**Fig. 3.**
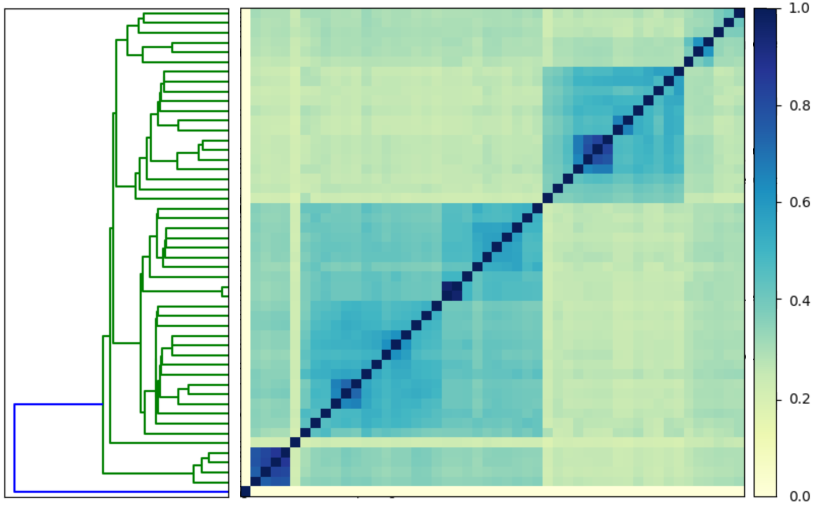
Heatmap and dendrogram generated using sourmash signatures built from 50 genomes that contained the word “*Escherichia coli*.” One signature is an outlier (highlighted in blue on the dendrogram).

**Fig. 4.**
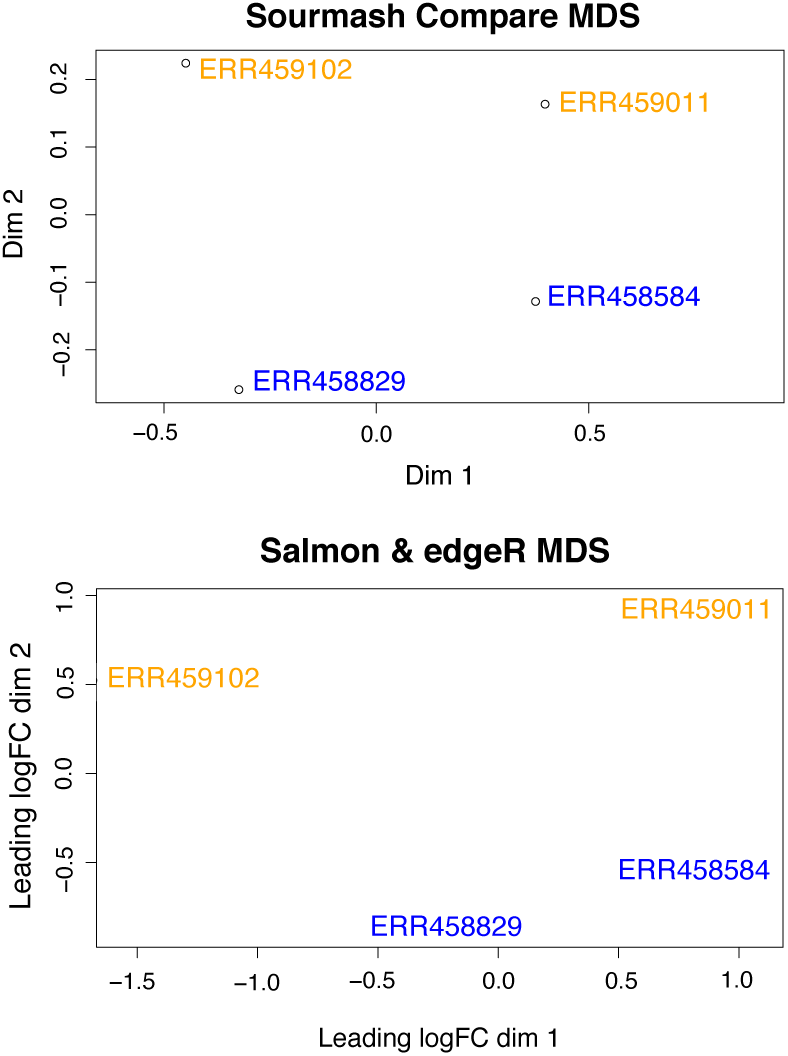
MDS plots produced via sourmash comparison of read data and via transcript quantification analysis are similar. Wild-type *S. cerevisiae* samples (ERR459011, ERR459102) are in yellow and mutant samples (ERR458584, ERR458829) in blue.

### Classifying signatures

The search and gather functions allow the user to classify the contents of a signature by comparing it to a database of signatures. Prepared databases for microbial genomes in RefSeq and GenBank are available at https://sourmash.readthedocs.io/en/latest/databases.html.

However, it is also simple to create a custom database with signatures.

Below we make a database that contains 50 *Escherichia coli* genomes.

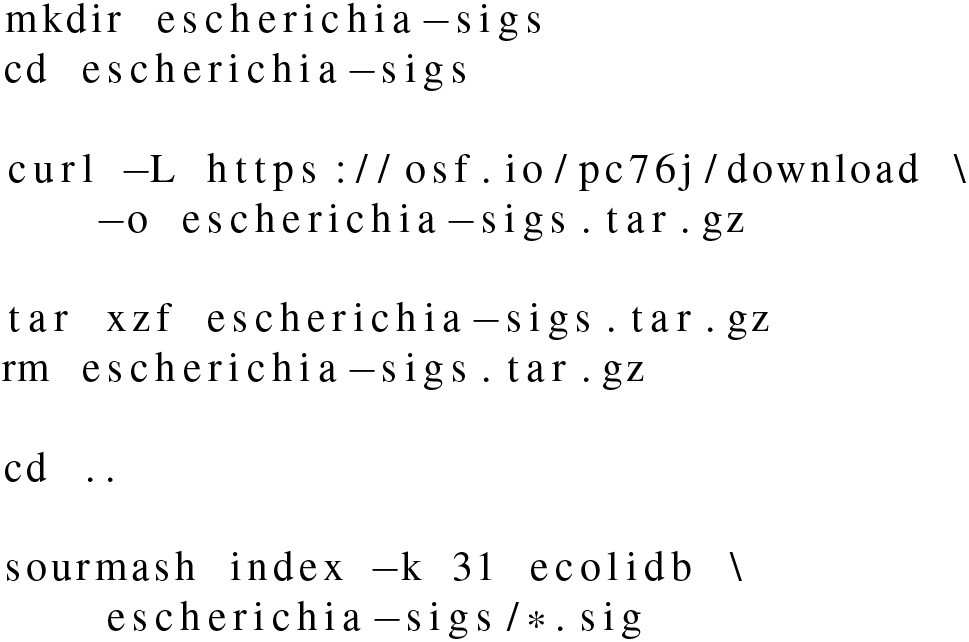

This database can be queried with search and gather using any signature calculated with a k-size of 31.

For example, below we download a small set of k-mer trimmed *Escherichia coli* reads and generate a signature with k=31.

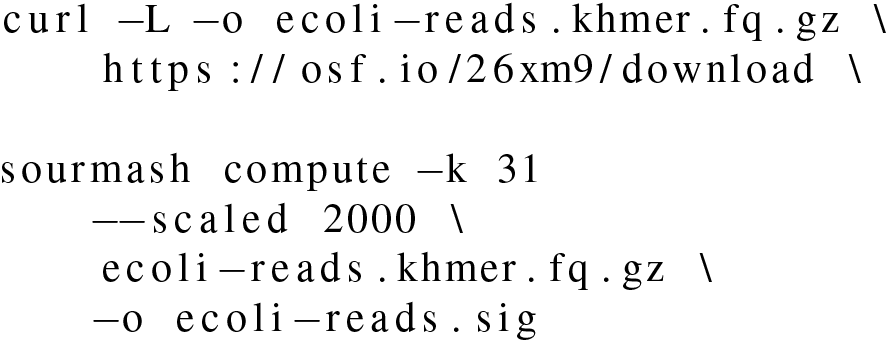

Then, we search the 50-genome database created above. sourmash search:

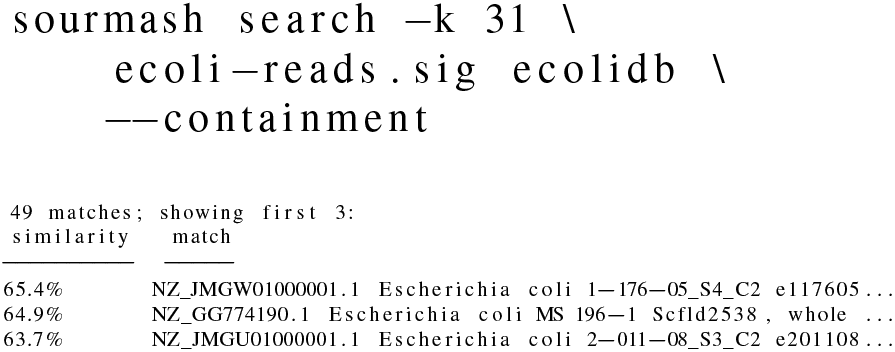

Breadth-first sourmash search finds all matches in the SBTMH that are present in the query signature (above a specified threshold).

Now try the same search using sourmash gather.

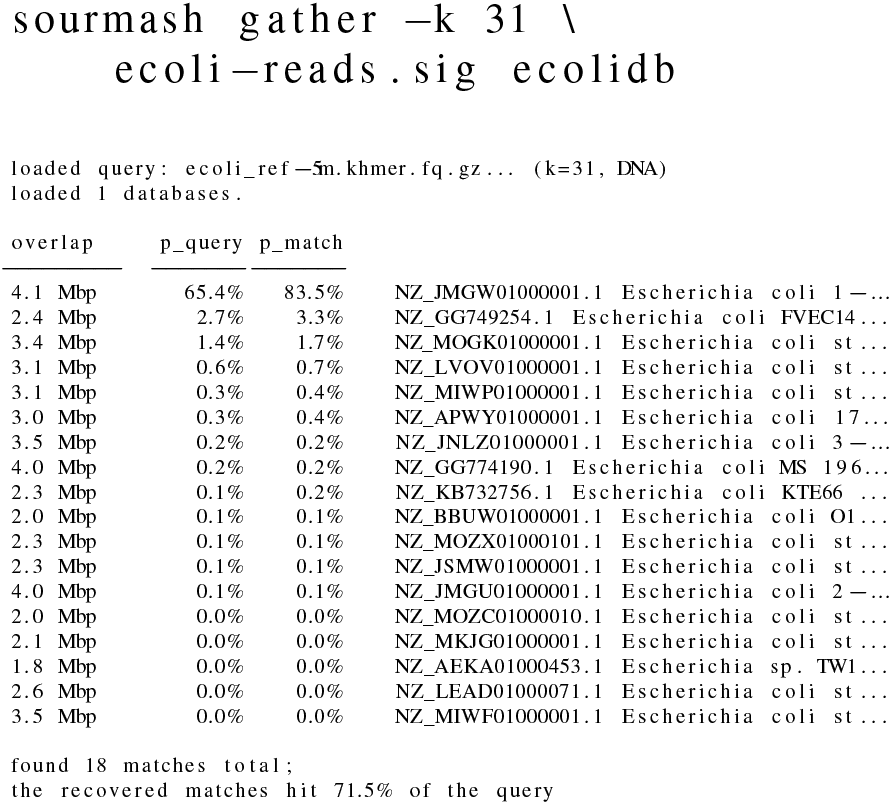

Best-first sourmash gather finds the best match first, e.g. here the first *E. coli* genome has an 83% match to 65.4% of our query signature. The hashes that matched (65.4% of the query) are then subtracted, and the database is queried with the remaining hashes (34.6% of original query). This process is repeated until the threshold is reached. sourmash postprocesses similarity statistics after the search such that it reports percent identity and unique identity for each match. sourmash gather is also useful for rapid metagenome decomposition. Below we calculate a signature of a metagenome using raw reads, and then use gather to perform a best-first search against all microbial genomes in Genbank. This approach was recently used to classify unknown genomes in a “mock” metagenome (21). The mock community was made to contain 64 genomes, but additional genomic material was inadvertently added prior to sequencing. Below we will use gather to investigate the content in the mock metagenome that did not map to the 64 reference genomes. For details on how this signature was created, please see (22). Note that the GenBank database is approximately 7.8 Gb compressed, and 50 Gb decompressed. Searches of the current GenBank database run fastest if allowed to use 11 Gb of RAM.

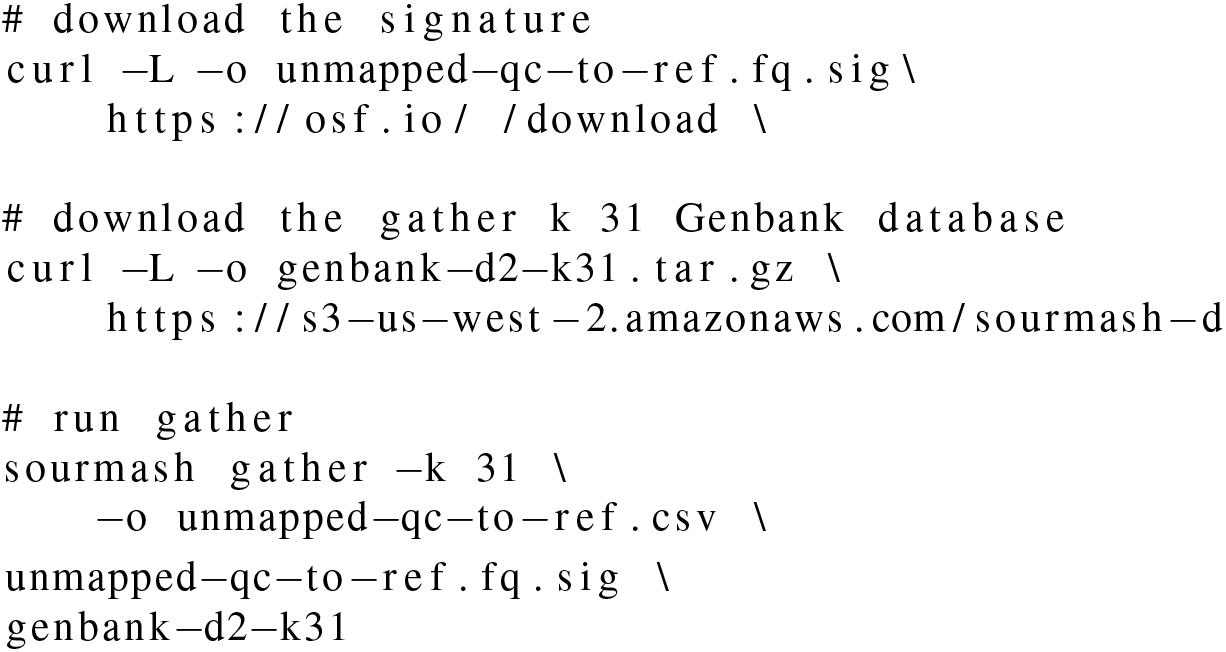

The output to the terminal begins:

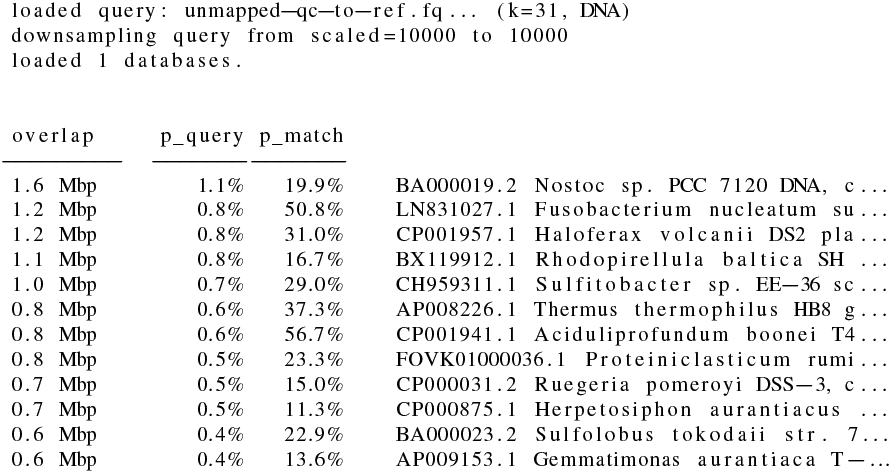

We see that 20.1% of k-mers match 82 genomes in GenBank. The majority of matches are to genomes present in the mock community. However, some species like *Proteiniclasticum ruminis* were not members of the mock community. These results also highlight how sourmash gather behaves with inexact matches such as strain variants. For example, we see two matches between *P. ruminis* strains among all matches. This likely indicates that a *P. ruminis* strain that has not been sequenced before is in our sample, and that it shares more k-mers of size 31 in common with one strain than the other. (See (23) for further analysis of this strain.)

sourmash gather and search also support custom databases. Using a custom database with sourmash gather, we can idnetify the dominant contamination in the domesticated olive genome (20). Below, we will use a database containing all fungal genomes in NCBI. We will then use the streaming compatibility of sourmash to download and calculate the signature. Lastly, we will search the olive genome against the fungal genomes using gather.

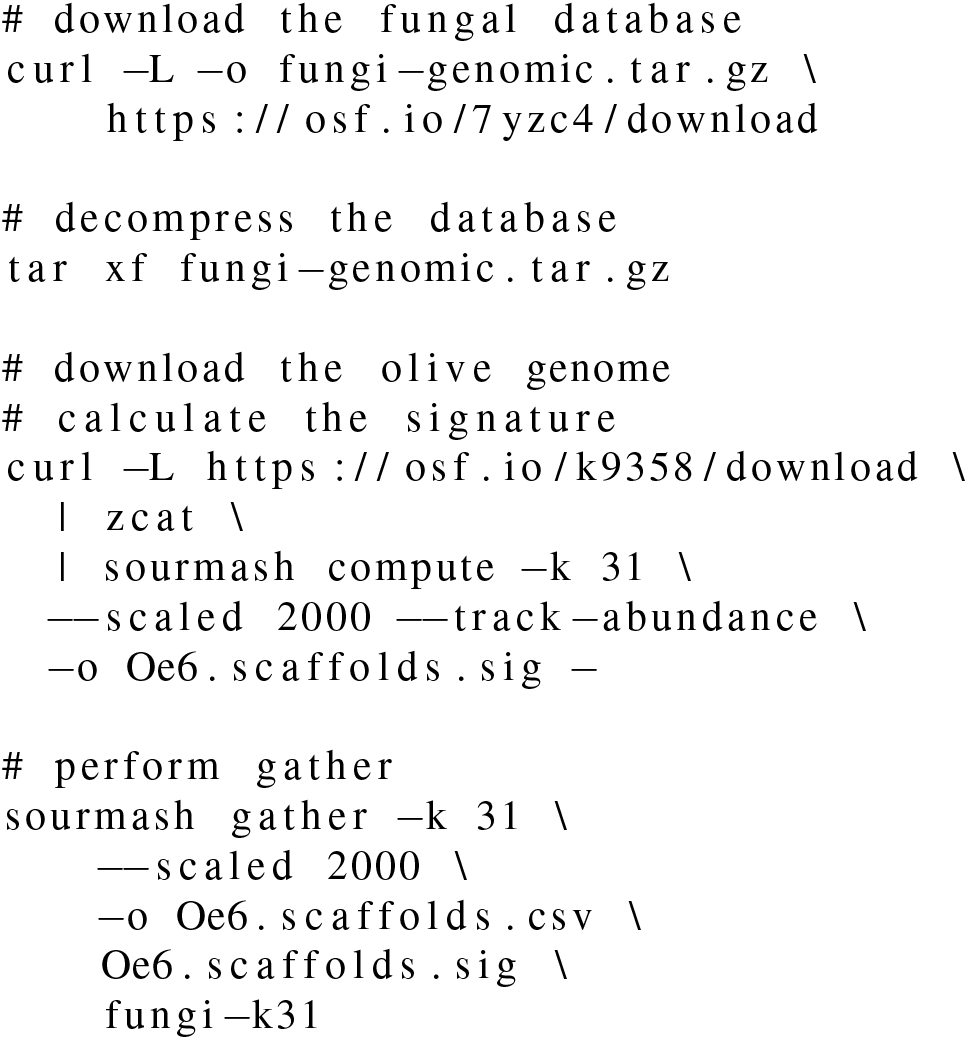

Using gather, we see two matches both within the genus *Aureobasidium*. This is the dominant contaminant within the genome (20).

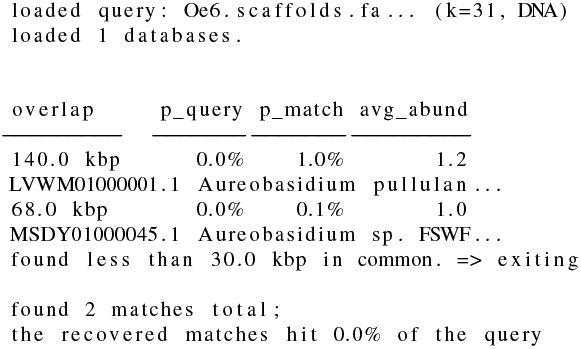

## Conclusions

The sourmash package provides a collection of tools to conduct sequence comparisons and taxonomic classification, and makes comparison against large-scale databases such as GenBank and SRA tractable on laptops. sourmash signatures are small and irreversible, which means they can be used to facilitate pre-publication data sharing that may help improve classification databases and facilitate comparisons among similar datasets.

## Author contributions

LI and CTB are the primary authors of the sourmash package. NTP, TR and CTB drafted the manuscript. NTP, TR, PB, LI, and CTB edited the manuscript and contributed to sourmash code, documentation, and use case development.

## Appendix

### MDS Comparison

Code to generate this MDS plot via Salmon (16) and edgeR (17) is available online at https://osf.io/97rt4/.

